# Anchoring the human olfactory system to a functional gradient

**DOI:** 10.1101/2020.03.19.998849

**Authors:** Alice Waymel, Patrick Friedrich, Pierre-Antoine Bastian, Stephanie J. Forkel, Michel Thiebaut de Schotten

## Abstract

Margulies et al. (2016) demonstrated the existence of at least five independent functional connectivity gradients in the human brain. However, it is unclear how these functional gradients might link to anatomy. The dual origin theory proposes that differences in cortical cytoarchitecture originate from two trends of progressive differentiation between the different layers of the cortex, referred to as the hippocampocentric and olfactocentric systems. When conceptualising the functional connectivity gradients within the evolutionary framework of the Dual Origin theory, the first gradient likely represents the hippocampocentric system anatomically. Here we expand on this concept and demonstrate that the fifth gradient likely links to the olfactocentric system. We describe the anatomy of the latter as well as the evidence to support this hypothesis. Together, the first and fifth gradients might help to model the Dual Origin theory of the human brain and inform brain models and pathologies.

---

Functional connectivity derived from functional magnetic resonance imaging relies on the low-frequency synchronisation of distant brain areas. Given that Blood-oxygen-level-dependent signal (BOLD) is a surrogate measure of neuronal activity (Ogawa et al., 1990) and because cells that fire together wire together (i.e neurophysiological rule, Hebb, 1949) it is assumed that the synchronisation of BOLD is a proxy of the connection between brain areas. And indeed, computational modelling indicated that resting state functional magnetic resonance imaging connectivity is partially constrained by the existence of direct and indirect structural connections (Honey et al., 2009).

Recently it has been demonstrated that functional connectivity can be reduced to distinct gradients (Coifman et al., 2005; Margulies et al., 2016) which summarise specific connectivity signatures distributed across the human brain. A recent study identified a principal gradient of functional connectivity which spans between primary sensory areas and the default mode network (Margulies et al., 2016) and may have shed light on evolutionary trends in the brain.

The dual origin theory (Abbie, 1940, 1942; Dart, 1934; Sanides, 1962) is an evolutionary hypothesis. The governing principle of this theory proposes that a progressive differentiation between the different layers of the cortex drives the difference in cortical cytoarchitecture between brain areas. This principle of cortical organisation is assumed to give rise to two functionally distinct systems respectively emerging from paleo- and archi-cortices. One system originates from the pyriform cortex and expands laterally towards the amygdala, the temporal, the orbitofrontal cortices and inferior parietal lobule and is referred to as the olfactocentric system. The other system arises from the hippocampus and progressively radiates medially along the cingulate cortex to reach the dorsolateral frontal cortex and superior parietal lobule, this system is referred to as hippocampocentric (for a comprehensive review on the dual origin see Pandya et al., 2017). Introducing functional gradients into this framework, the principal gradient seems to be represented by the hippocampocentric system (for anatomical reviews see Alves et al., 2019; Catani et al., 2013). This system is crucial for limbic and transmodal cognitive functions (e.g. episodic memory). Here, we hypothesise that one of the four additional gradients described by Margulies et al. (2016) should correspond to the second olfactocentric system.

## Methods

We downloaded the latest version of the five functional connectivity gradients available online (https://neurovault.org/collections/1598/). Meta-analytic maps of functional activations related to ‘olfactory’ (74 studies) and ‘episodic memory’ (332 studies) were harvested from neurosynth (https://neurosynth.org).

Next, the five gradients and the metanalytic maps were characterised using the multimodal parcellation of human cerebral cortex (Glasser et al., 2016). As subcortical areas also play an important role in cognition, we added 12 additional regions, defined manually, including the bilateral amygdala, caudate nucleus, hippocampus, pallidum, putamen, and thalami. We then created a matrix consisting of the average value for all voxels of each parcel (i.e. one parcel per row) for each gradient and the metanalytic maps (i.e. one map per column). Finally, similarities between gradients and metanalytic maps were assessed using bootstrapped (n = 1000 samples) Spearman rank correlation coefficient (Spearman, 1904) for non 0 values. We only report results significant after Bonferroni correction for multiple comparisons (n = 10 comparisons).

## Results

Meta-analytic activations maps from neurosynth indicate higher values for more replicable loci of activation. Accordingly, Spearman rank correlations confirmed that the higher the meta-analytic activations for episodic memory, the higher the value in the principal gradient (rho = 0.273, p < 0.001; figure 1). Similarly, correlations indicated that the higher the meta-analytic activations for olfaction, the higher the value of the fifth gradient (rho = 0.372, p < 0.001). No other correlation survived Bonferroni correction for multiple comparisons.

**Figure 1:**
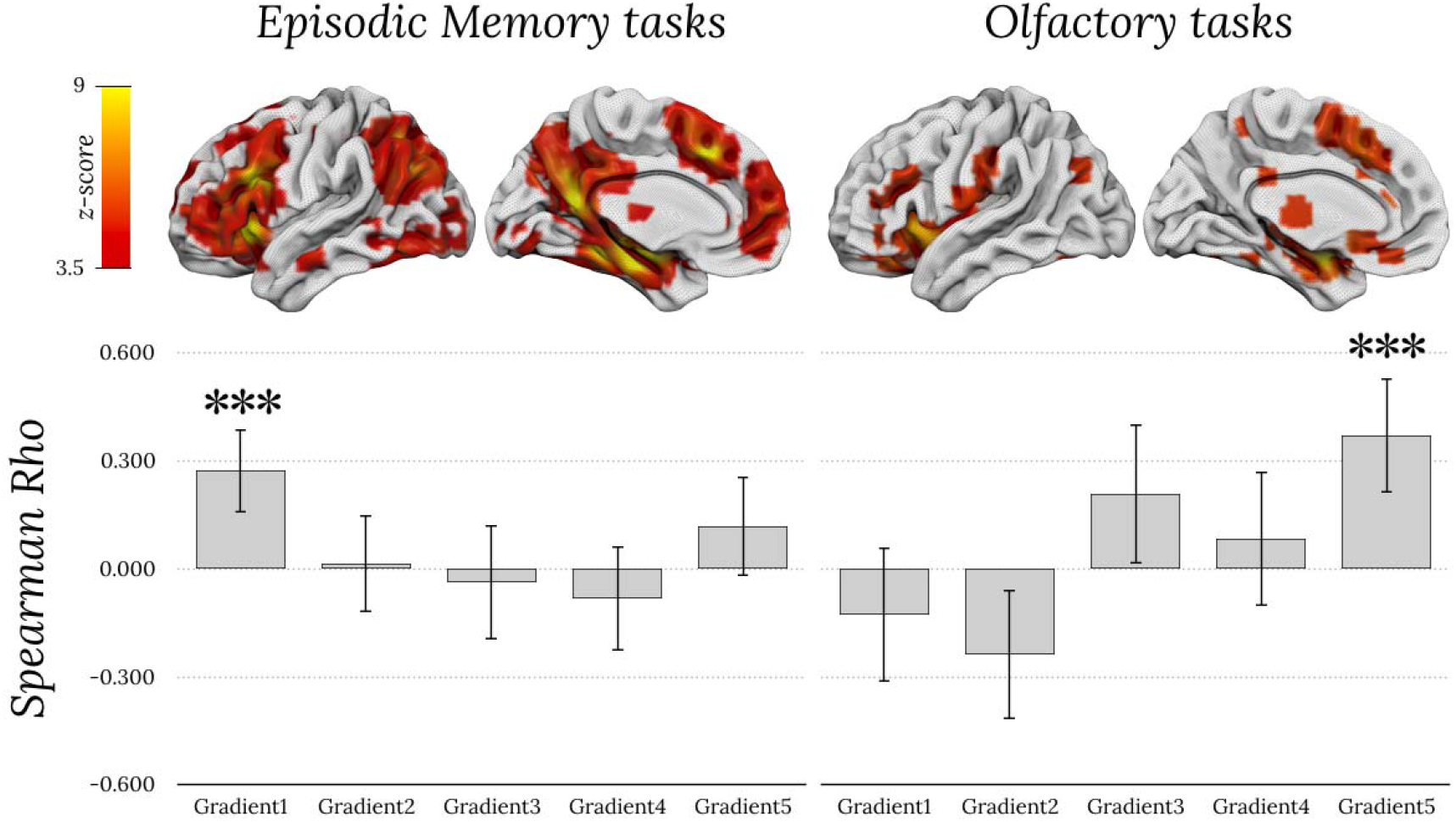
Spearman rank correlations between fMRI metanalytic maps of episodic memory tasks (left), olfactory tasks (right), and values of the five gradients. Top row corresponds to meta-analytic activations maps from neurosynth. Error bars corresponds to 95% confidence intervals. *** indicates p < 0.001 and is significant after Bonferroni correction for multiple comparisons.

Our statistical exploration of these gradients indicates that there is a previously undescribed link between the fifth gradient and the olfactocentric system, centred around the piriform cortex. Anatomically, Figure 2 confirms that the positive end of the gradient anchors to cortical and subcortical olfactory structures, whilst the negative end of the gradient includes brain regions unrelated to olfactory processing. At the cortical level, the positive end of the gradient includes the piriform, orbitofrontal, entorhinal, anterior insular, cingulate, and inferior temporal cortices, as well as the temporal pole, the posterior inferior frontal and the anterior supramarginal gyrus. At the subcortical level, the amygdala and the hippocampus are primarily involved.

**Figure 2:**
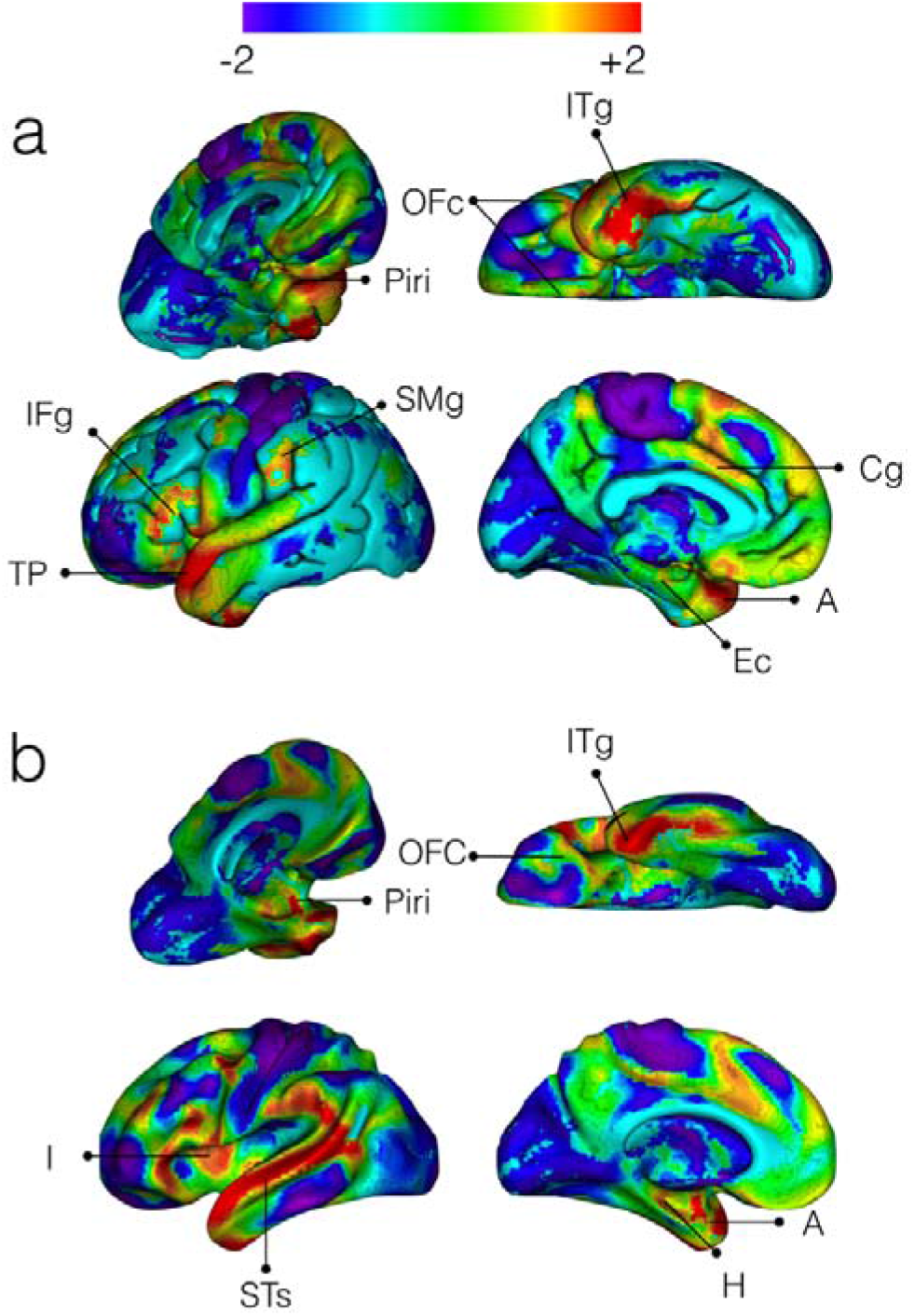
The fifth gradient of functional connectivity in the human brain. a) Shows the cortical surface representation of the gradient while b) demonstrates the inflated white matter surface. *ITg: Inferior Temporal gyrus; OFc: Orbitofrontal cortex; Piri: Piriform cortex; IFg: Inferior Frontal gyrus; TP: temporal pole; SMg: Supramarginal gyrus; Cg: Cingulate gyrus; I: Insula; STs: Superior Temporal sulcus; Ec: Entorhinal cortex; A: Amygdala; H: Hippocampus*

## Discussion

This study investigated the relationship between the previously reported five independent functional connectivity gradients in the human brain and the two system of the Dual Origin theory. Results are twofold. First, we confirmed the spatial correspondence between the first gradient and the hippocampocentric division. Second, we discovered a relationship between the positive end of the fifth gradient and olfactory activations. These results have implications for the study of the Dual Origin theory.

The expected relationship between the first gradient and episodic memory activation confirms the correspondence of the principal gradient with the hippocampocentric system of the Dual Origin theory. According to the theory, the hippocampocentric system should link previously learned information with sensory information (Pandya and Yeterian, 1990). Similarly the principal gradient links episodic memories centers to primary sensory motor cortices. The correspondence between the first gradient and the hippocampocentric system suggests that functional gradients might represent some of the variance related to the progressive differentiation of the cortical layers proposed by the Dual Origin theory. If this assumption is true, another gradient should reflect the olfactocentric division.

The gradient that anatomically is most closely associated with the olfactory system is the fifth gradient. The fifth gradient involves a set of regions that likely corresponds to the olfactocentric system and is classically activated in *f*MRI studies during olfactory-specific tasks. The olfactocentric system is traditionally divided into the primary olfactory cortex and the secondary olfactory cortex (Purves and Williams, 2001) with a high modularity (Arnold et al., 2019). The primary olfactory cortex is represented by the piriform cortex that projects to various cortical (e.g. orbitofrontal, entorhinal, and insular cortex) and subcortical structures (e.g. amygdala, hypothalamus). The secondary olfactory cortex comprises the orbitofrontal, insular, and cingulate cortices and the hippocampus subcortically. Structures within both these olfactory systems and some of their projections are included in the fifth gradient as shown by Margulies et al. (2016). The hypothalamus is not included in the functional connectivity gradient, although it also belongs to the olfactocentric system. As one limitation of surface analyses is that they notoriously underestimate deep subcortical structures, this may might explain this discrepancy (Alves et al., 2019).

The primary and secondary olfactory system are selectively recruited according to task demands. For odour detection tasks, the piriform cortex is activated, whereas for familiarity judgements, the orbitofrontal cortex is recruited (Royet et al., 2001). There are also amygdala and cingulate cortex activations during judgements of the pleasantness of odours (Savic et al., 2000). The olfactory system also links with hippocampocentric centres including the hippocampus and entorhinal cortex, both essential structures supporting memory (Corkin, 2013). Indeed, odours are well known to be particularly powerful memory triggers (Chu and Downes, 2002; Herz and Cupchik, 1995). As such, the olfactory memory system is unique in that it does not entail thalamic relays and can facilitates fast access to very old and specific episodic memories (Willander and Larsson, 2006). Hence, the link between the piriform cortex, hippocampus, and entorhinal cortex might serve as a powerful complementary memory system, while other areas may flavour olfactory perception with intensity and emotion.

The fifth gradient also includes additional regions that are not strictly part of - but can be associated with - the olfactocentric system, such as the temporal pole and superior temporal sulcus, posterior ventral prefrontal cortex, supramarginal gyrus and pre-Supplementary Motor Area (preSMA). Even though these structures fall outside the definition of the olfactory system, the temporal pole and superior temporal sulcus are closely related to the semantic memory system (Chiou and Lambon Ralph, 2016; Mesulam et al., 2015) and might be necessary for the semantic labelling of odours. Similarly, the posterior ventral prefrontal cortex, supramarginal gyrus, insula and preSMA together resemble the saliency network, which is critical for detecting behaviourally relevant stimuli (Corbetta and Shulman, 2002) which might extend to olfaction. As previously reported (Aboitiz and Montiel, 2015), a link between odour, attention and behaviour seems to be intuitively plausible for survival and could thus have been phylogenetically relevant. Hence, albeit not strictly part of the olfactory system as we know it, these additional regions might be consistent with an extended system linking olfactory perception, semantic memory, and awareness (i.e. saliency).

Linking the fifth gradient to the olfactocentric system nicely complements the first gradient described by Margulies et al. (2016) in light of the dual origin theory (Dart, 1934; Pandya et al., 2017; Sanides, 1964). However, one might notice that the first and fifth gradient were not equally represented across the human brain. Several interpretations are available to explain this difference. First, the olfactory system (e.g. temporal pole, orbitofrontal cortex) is largely located near the noisiest area of the resting-state fMRI imaging (Olman et al., 2009) which might have compromised the variance decomposition of the signal and may, therefore, account for a smaller percentage of variance. The olfactocentric division might also be less prominently represented in humans compared to other species. Comparative anatomy studies suggest that, with evolution, olfactory structures have increased in absolute size much less than the cortical surface (Reep et al., 2007). This means that the olfactory system would be smaller in the human brain by comparison. Another aspect that may explain a smaller olfactocentric division could be the easier access to food sources in humans. In accordance with the Ecological Brain hypothesis (Milton, 1981), the relative size of the olfactory system (Jacobs, 2012) and the orbitofrontal cortex (Louail et al., 2019) would correlate with the ability to identify access to food resources. Given that humans have easy access to food since birth, the functional connectivity within this system might have been partially pruned during brain development or through evolution. The availability of new open datasets for comparative research on the functional connectivity of primate species (Milham et al., 2018; Thiebaut de Schotten et al., 2019; Workshop et al., 2020) might confirm the hypothesis that the olfactocentric division is relatively smaller in humans compared to other primates. Finally, it is also possible that the archicortex (i.e. olfactrocentric) might have left an evolutionary signature different from the paleocortex (i.e. hippocampocentric) and that the archicortex’s signature is not as well visible with functional connectivity. Promising research exploring the structural rather than functional primate connectome may shed light onto this hypothesis (Blazquez Freches et al., in press; Mars et al., 2018; Mars et al., 2016).

Overall, the potential characterisation of the hippocampocentric and olfactocentric systems via the first and fifth gradients might shed light onto phylogenetic principles of functional brain organisation. This characterisation may provide a new tool to scrutinise hypotheses about evolutionary and developmental changes of the brain as well as stratify patterns of neurodegenerative disease progression along the hippocampocentric or olfactocentric systems.

## Acknowledgments

This project has received funding from the European Research Council (ERC) under the European Union’s Horizon 2020 research and innovation programme (grant agreement No. 818521).

